# Crosstalk between Flavonoids and the Plant Circadian Clock

**DOI:** 10.1101/2021.07.15.452546

**Authors:** Sherry B. Hildreth, Evan S. Littleton, Leor C. Clark, Gabrielle C. Puller, Shihoko Kojima, Brenda S.J. Winkel

**Affiliations:** Department of Biological Sciences, Virginia Tech, Blacksburg, VA 24061; Fralin Life Sciences Institute, Virginia Tech, Blacksburg, VA 24061; Molecular and Cellular Biology graduate program, Virginia Tech, Blacksburg, VA 24061; Mass Spectrometry Incubator, Virginia Tech, Blacksburg, VA 24061

## Abstract

Flavonoids are a well-known class of specialized metabolites that play key roles in plant development, reproduction, and survival. Flavonoids are also of considerable interest from the perspective of human health, both as phytonutrients and pharmaceuticals. RNA-Seq analysis of an Arabidopsis null allele for chalcone synthase (CHS), which catalyzes the first step in flavonoid biosynthesis, has uncovered evidence that these compounds influence the expression of circadian clock genes in plants. Analysis of promoter-luciferase constructs showed that the transcriptional activity of genes encoding two components of the central clock, *CCA1* and *TOC1*, across the day/night cycle is altered in CHS-deficient seedlings. The effect of flavonoids on circadian function was furthermore reflected in photosynthetic activity, with chlorophyll cycling abolished in the mutant line. Analysis of a mutant lacking flavonoid 3’-hydroxylase (F3’H) activity, and thus able to synthesize mono- but not di-hydroxylated B-ring flavonoids, suggests that the latter are at least partially responsible, as further supported by the effects of quercetin on *CCA1* promoter activity in wild-type seedlings. Collectively, these experiments point to a previously-unknown connection between flavonoids and circadian cycling in plants and open the way to better understanding of the molecular basis of flavonoid action.

## Introduction

Plants produce an astonishing array of natural products via complex networks of specialized metabolism (Fernie and Tohge, 2017). These unique biochemicals, which number in the hundreds of thousands across the plant kingdom, are essential for plant growth, survival, and reproduction, and are critical mediators of interactions with other organisms. Flavonoids are one of the major groups of specialized metabolites that are present in virtually every member of the plant kingdom. The biosynthetic pathway present in extant plant species has been traced back to the emergence onto land some 550-470 million years ago and to have been key to the eventual colonization of virtually every terrestrial niche (reviewed in Yonekura-Sakakibara et al., 2019; Davies et al., 2020). It has been hypothesized that flavonoids initially functioned as physiological regulators and chemical messengers, consistent with the presumed very low concentrations and simple structures produced by the ancestral pathway (Stafford, 1991). This system evolved to enable the production of the multiple classes and more than 9,000 different structures documented to date with diverse roles that include mediating phytohormone activity and reproduction, defense from predators and pathogens, and protection from heavy metals, UV light, and other stresses. Flavonoids also play key roles in positive interactions with other organisms, including in recruiting pollinators, modulating the microbiome, and conversely, serving as abundant phytonutrients in the human diet (Crozier et al., 2009; Wang et al., 2018) and of long-standing interest as pharmaceuticals (Guo et al., 2019; Maher, 2019).

Plant flavonoid biosynthesis is mediated by highly-conserved machinery, with the first two steps catalyzed by enzymes derived from fatty acid metabolism and subsequent reactions by members of more broadly-dispersed enzyme superfamilies (Yonekura-Sakakibara et al., 2019). This system is organized as a multienzyme complex, which in at least some cases appears to be assembled around the entry-point enzyme, the type III polyketide synthase chalcone synthase (CHS) (Nakayama et al., 2019), with competitive interactions controlling the distribution of flux into branch pathways (Crosby et al., 2011). This enzyme complex is localized, in part, at the cytoplasmic face of the endoplasmic reticulum, with the resulting end products (flavonols, anthocyanins, and proanthocyanidins in Arabidopsis) transported primarily to the vacuole and cell wall (Agati et al., 2012; Zhao, 2015). Flavonoid enzymes have also been reported to localize to the nucleus in numerous species, where they may be responsible for the synthesis of flavonoids *in situ* (Winkel, 2019; Zhang et al., 2020). These proteins may also have alternative functions, as suggested by our recent report that CHS interacts with MOS9, a nuclear protein required for defense gene expression with no known function in flavonoid metabolism (Watkinson et al., 2018).

To explore CHS function from a broader perspective, RNA-Seq was used to examine the effects of a null mutation in Arabidopsis CHS (*tt4-11*; Figure 1A) on global gene expression. Surprisingly, this analysis uncovered alterations in the expression of the central circadian clock genes as well as a large number of clock-regulated genes. These findings were corroborated in complementary experiments, including showing effects at the level of transcriptional control of the core clock genes, *CLOCK ASSOCIATED 1* (*CCA1*) and *TIMING OF CAB EXPRESSION 1* (*TOC1*), as well as disruption of cycling of photosynthetic activity, a key indicator of endogenous circadian rhythms. Analysis of a *tt7* mutant line, which is deficient in flavonoid 3’-hydroxylase (F3’H) activity (Figure 1A), suggests that dihydroxylated flavonoids may be responsible for the observed effects of flavonoids on clock function. Collectively, the data point to a previously-unknown function for these ubiquitous plant metabolites.

**Figure 1.**
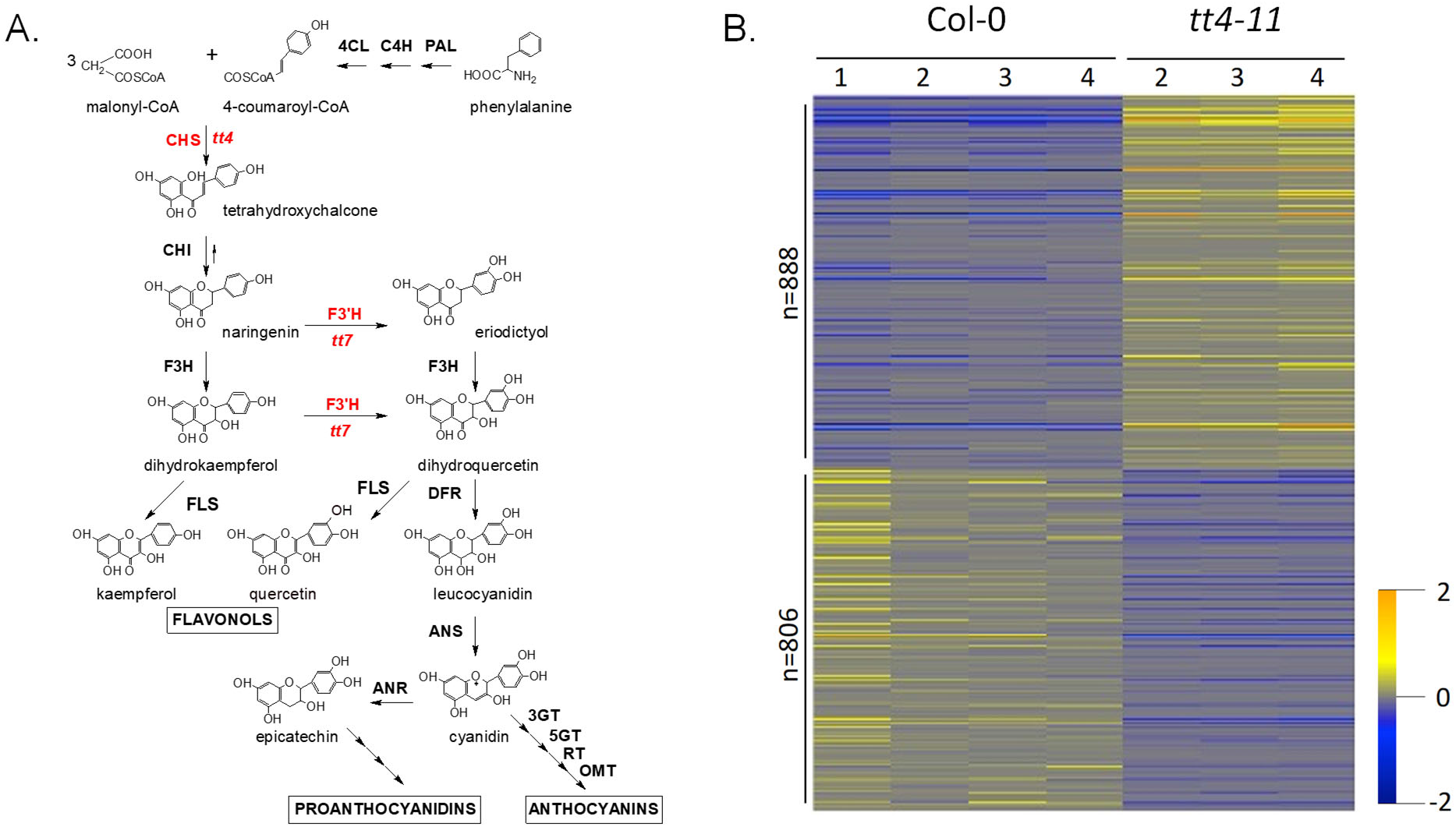
Effects of disruption of flavonoid metabolism on the seedling transcriptome. A. Simplified schematic of the Arabidopsis flavonoid pathway showing locations of the enzymatic steps disrupted in *tt4-11* and *tt7-5*. The two enzymes and corresponding loci that are the focus of this study are highlighted in red text. Boxed text identifies the three major flavonoid end products in Arabidopsis. Enzyme names are abbreviated as follows: phenylalanine ammonia-lyase (PAL), chalcone synthase (CHS), chalcone isomerase (CHI), flavanone 3-hydroxylase (F3H), flavonoid 3’-hydroxylase (F3’H), dihydroflavonol 4-reductase (DFR), anthocyanidin synthase (ANS), 3- and 5-glucosyl transferase (3GT and 5GT),rhamnosyl transferase (RT), and *O*-methyl transferase (OMT). B. Heatmap of z-scores for DEGs meeting the criteria log2fold change ≤ 0.50 and p ≤ 0.05. Color scale: yellow = elevated and blue = reduced in *tt4-11*.

## Results

### CHS-deficient lines exhibit altered expression of numerous clock-associated genes

While the primary site of flavonoid metabolism in cells has long been held to be the cytoplasmic face of the endoplasmic reticulum, a number of studies in recent years point to the nucleus as another site of active flavonoid biosynthesis as well as potential moonlighting roles for the enzymes (Winkel, 2019; Zhang et al., 2020). This raises the possibility that flavonoid enzymes or end products may directly or indirectly influence gene expression. To examine this further, we undertook transcriptome analysis of a well-characterized Arabidopsis flavonoid null mutant, *tt4-11*. This allele carries a T-DNA insertion in the second exon of the CHS gene, resulting in loss of all detectable CHS protein and flavonoid end products (Lewis et al., 2011; Bowerman et al., 2012). RNA-Seq was used to compare the transcriptomes of Col-0 wild type and *tt4-11* seedlings grown under conditions of maximal flavonoid accumulation, consisting of six days of germination on 1X MS; 2% sucrose under continuous white light illumination (LL) (Dataset S1). This analysis revealed just 163 genes with differences in expression meeting a stringent criterion of log2fold ≥ |±1.0| (= ±2-fold change), all with p values <<0.001 (Dataset S2). Of these differentially-expressed genes (DEGs), 121 had elevated transcript levels in *tt4-11* and 42 (besides *CHS*) had levels that were reduced. Many more genes met the more conventional cutoff of log2fold |±0.5| (~1.4 fold) and p≤0.05, 1694 total, with 888 upregulated and 806 downregulated (not including CHS) in the mutant line (Figure 1B; Dataset S3).

The 163 high-confidence DEGs encompassed several functional groups containing multiple members. These included five ubiquitin ligases and eleven genes involved in ethylene response or metabolism, all with elevated levels in *tt4-11* (2.0 to 3.2-fold), as well as a number of transporters and glycine-rich RNA binding proteins that exhibited reduced transcript levels (2.0 to 3.7-fold change). Unexpectedly, one of the largest functional groups encoded components of the core circadian clock and clock-associated proteins, a total of ten of the 163 high-confidence DEGs (Table 1). This was also reflected in the PANTHER GO-Slim Biological Process analysis (Mi et al., 2020), with the highest enrichment (44.29 fold) for genes associated with circadian rhythms (Dataset S4). In fact, the clock-associated gene, *CCL,* had the most elevated transcript levels of any gene in the dataset (log2fold 1.81 = 3.5 fold), while transcripts for the key circadian transcriptional repressor, *CCA1*, were the third most reduced (log2fold 2.13 = 4.4 fold). The high-confidence DEGs also exhibited a substantial (almost 2-fold) enrichment of predicted circadian-regulated genes, with 60% belonging to this group in *tt4-11* (64 of 106 expressed genes) (Dataset S2) compared to just 32% for the wild-type circadian transcriptome (Covington et al., 2008). Consistent with this finding, a large number of the high-confidence DEGs contained CCA1 binding sites previously identified by ChIP-Seq (Dataset S3; Nagel et al., 2015): 16 (13%) of those with elevated transcript levels in *tt4-11* and 27 (64 %) with reduced levels. It is also worth noting that members of the functional classes mentioned above have associations with clock control: two of the five ubiquitin ligases with elevated levels in *tt4-11* are among those identified a potential regulators of circadian function by Feke et al. (2019); ethylene is key to regulation of the clock in response to biotic factors, which in some cases involves specific ethylene response factors (ERFs) (Haydon et al., 2017; Li et al., 2019); and one member of the glycine-rich RNA-binding protein group, AtGRP7, has been shown to regulate the circadian cycling of its own transcript (Staiger and Heintzen, 1999). Many more core clock and clock-associated transcripts met the more conventional log2fold |±0.5| (~1.4 fold) cutoff with p≤0.05, including virtually all components of the Arabidopsis clock machinery (Singh and Mas, 2018). This larger group of DEGs was also enriched for predicted circadian-regulated genes (46% compared to 32% for all genes in the Covington/Edwards collection; Dataset S3), although not quite as much as the high-confidence DEGs (at 60%).

**Table 1:** Clock genes with altered transcript levels in CHS/ flavonoid-deficient *Arabidopsis* plants relative to wild type^1^.

| Gene Symbol | Gene name | log2Fold Change | p-value |
| --- | --- | --- | --- |
| **CORE CLOCK** |  |  |  |
| CCA1 | AT2G46830.1 | 2.1283 | 6.96E-28 |
| CCA1 | AT2G46830.2 | 1.5956 | 7.66E-10 |
| LNK2 | AT3G54500.3 | 1.2121 | 2.29E-06 |
| RVE8 | AT3G09600.1 | 1.0339 | 6.00E-05 |
| RVE2 | AT5G37260.1 | 0.3685 | 0.0200 |
| ELF3 | AT2G25930.1 | -0.3349 | 0.0323 |
| PRR7 | AT5G02810.1 | -0.4250 | 0.0196 |
| TOC1 | AT5G61380.1 | -0.4918 | 1.79E-04 |
| PRR5 | AT5G24470.1 | -0.7037 | 2.52E-03 |
| BOA | AT5G59570.1 | -0.8615 | 8.76E-04 |
| ELF4 | AT2G40080.1 | -1.1359 | 6.44E-11 |
| **CLOCK-ASSOCIATED** |  |  |  |
| LNK3 | AT3G12320.1 | 1.4038 | 1.53E-08 |
| REV3 | AT1G01520.1 | 1.1400 | 2.81E-06 |
| CCL1 | AT5G65480.1 | 0.3994 | 5.64E-03 |
| CRY1 | AT4G08920.1 | 0.3248 | 5.90E-03 |
| NUP205 | AT5G51200.1 | 0.3209 | 9.09E-03 |
| CKA2 | AT3G50000.1 | 0.2945 | 0.0141 |
| CKA1 | AT5G67380.1 | -0.6177 | 0.0361 |
| ARR4 | AT1G10470.1 | -0.7879 | 1.03E-07 |
| GI/FB | AT1G22770 | -1.0197 | 6.50E-08 |
| ADO3 | AT1G68050.1 | -1.6202 | 1.89E-11 |
| CCL | AT3G26740.1 | -1.8140 | 9.35E-36 |

<sup>1</sup> Positive values indicate elevated expression in *tt4-11* relative to wild type. Color scale: red = elevated, blue = reduced in *tt4-11*. Genes not meeting the log2fold $\geq |0.5|$ cutoff are shown in gray font.

To validate the results of the transcriptome experiment and further explore these initial findings, qPCR was used to compare the expression of several of the clock-associated genes under both LL and 12h/12h LD conditions. For this analysis, comparisons were made in *tt4-11* and Col-0 seedlings, as well as in another CHS null allele, *tt4-2(bc)* (Burbulis et al., 1996; Bennett et al., 2006). As illustrated in Figure 2, the differences detected by qPCR were smaller than observed by RNA-Seq and statistically significant in only some cases, due in part to the relatively low levels of expression of these genes, particularly in LL. Overall, however, the expression patterns in seedlings entrained to LL mirrored those observed in the transcriptome dataset in both *tt4* alleles, thereby also confirming that the observed effects are linked to the *CHS* locus. Surprisingly, the opposite effect on clock gene expression was observed in seedlings entrained to LD, at both the morning and evening timepoints (Figure 2B, AM and PM, respectively). The differential effects under LD and LL conditions may reflect a further role for flavonoids in the response to light stress that is overlaid on the contribution to circadian homeostasis.

**Figure 2.**
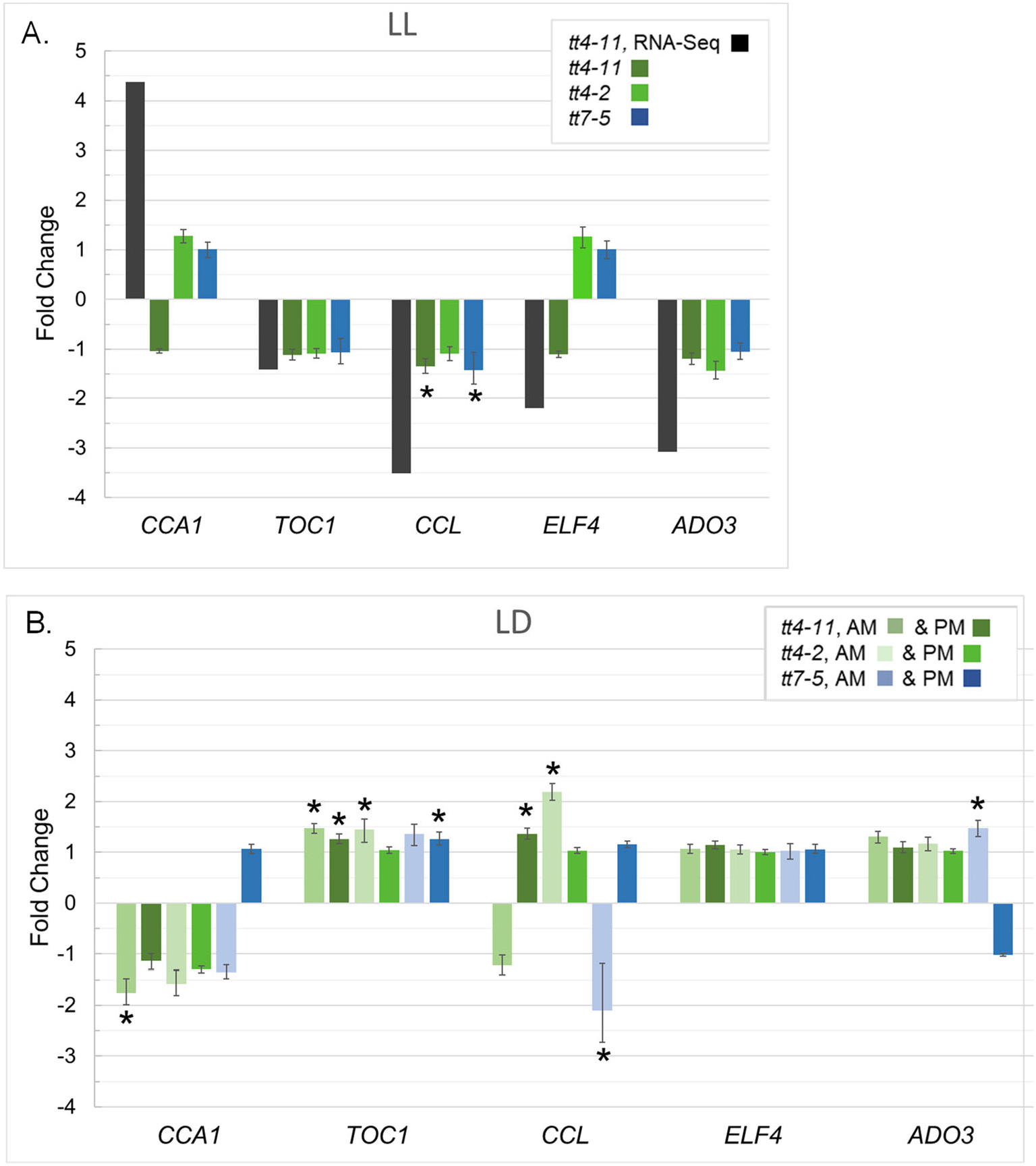
qRT-PCR analysis of select clock-associated genes in flavonoid mutant lines. Differences are shown as fold change in gene expression in mutant lines relative to the corresponding wild type; error bars are standard errors (n=3 biological replicates), with asterisks indicating a significant difference (p<0.05) from the corresponding wild type per Student’s t-test. (A) Seedlings grown in continuous light (LL); RNA-Seq data converted to fold change are shown for comparison. (B) Seedlings entrained in 12h/12h LD with samples collected at 4h (AM) and 12h (PM) after lights came on.

### *tt4* alleles alter *CCA1* and *TOC1* promoter activity *in vivo*

To determine whether the changes in steady state mRNA levels observed in *tt4* might originate at the transcriptional level, we took advantage of Arabidopsis lines containing the promoter-luciferase (LUC) constructs, *CCA1p::LUC* or *TOC1p::LUC* (Salomé and McClung, 2005). These lines have been used extensively to monitor circadian transcriptional activity in intact plants in real time. The two promoters exhibit complementary patterns of expression, with *CCA1* maximal in the morning and *TOC1* in the evening, approximately 12 hours out of phase. The two constructs were introduced into *tt4-11* by crossing with the wild-type parental lines, followed by identification of multiple independent F3 lines homozygous for *tt4-11 and* expressing the transgene. Seedlings were germinated on MS-sucrose plates for 5 days under 12h/12h LD conditions and then moved to constant darkness (DD) and LUC activity was monitored for approximately 10 days. The resulting traces showed that *CCA1* and *TOC1* promoter activity mirrored transcript levels (Figure 2B), with expression elevated and reduced, respectively, in *tt4-11* relative to wild type seedlings across the day/night cycle (Figure 3A).

**Figure 3.**
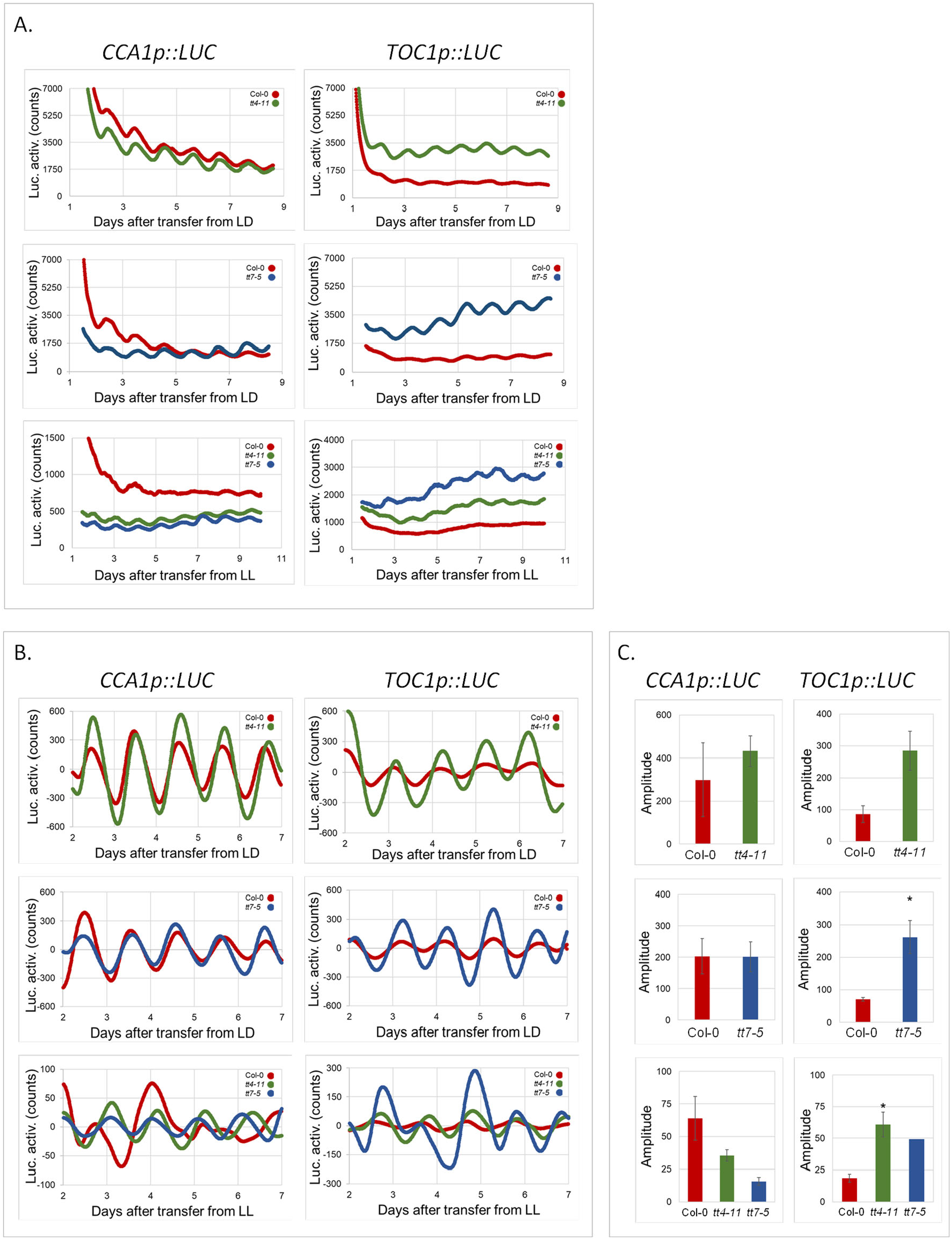
Effects of flavonoid mutations on *CCA1* and *TOC1* promoter activity. Luciferase activity was assessed in seedlings following transfer from 12h,12h LD (top two rows) or LL (bottom row) to continuous darkness (DD) using a Lumicycle luminometer. In all panels, results are shown for Col-0 (red), *tt4-11* (green), and *tt7-5* (blue) seedlings. Each line represents the average of four biological replicates for Col-0 and the average of 2-4 replicates for each of three independent F3 lines for *tt4-11* and *tt7-5*. (A) Unprocessed and (B) detrended (baseline-subtracted) data; (C) Comparison of amplitudes of detrended data; error bars are standard error; * indicates Student’s t-test p-value <0.05.

The data were detrended (baseline-subtracted) and statistical analysis of amplitude, phase, and period performed using BioDare2 (Figure 3B and C). This revealed significant changes in amplitude for *TOC1* in both mutant backgrounds, consistent with the role of this transcription factor in mediating clock function in response to a wide range of conditions, including abiotic factors such as drought (Legnaioli et al., 2009), temperature (Zhu et al., 2016), and light (Soy et al., 2016). There were also modest effects on period and phase in *tt4-11* (Figure S1). In contrast, little effect was observed on the amplitude, phase, or period of *CCA1* promoter activity in LD, (Figure S1), reflecting the results of a study by Pruneda-Paz et al. (2014) that failed to identify transcription factors mediating *CCA1* amplitude. Crosstalk between flavonoids and the plant clock under LD conditions does appear to occur, at least in part, at the level of transcription, particularly in the case of *TOC1*, as well as through post-transcriptional processes in the case of *CCA1*.

A parallel analysis performed with seedlings entrained to LL conditions showed that *CCA1* and *TOC1* promoter activity was substantially lower overall in LL than in LD (top and middle panels versus bottom panels, Figure 3A). This is consistent with the overall lower steady-state RNA levels observed by qPCR. However, relative promoter activities in the mutant and wild-type lines were the same as in LD, and therefore opposite the trends observed for steady-state transcripts, where the levels of *CCA1* were higher and *TOC1* lower in LL (Table 1, Figure 2). The same pattern was displayed in the relative amplitudes of the detrended traces, with *CCA1* having an apparent reduction and *TOC1* an apparent elevation, although only the latter was statistically significant due to the low and variable amplitudes of the rhythms in LL (Figure 3B and C). This suggests that, if flavonoids do play an additional role in seedlings under light stress, this may be achieved via post-transcriptional control.

### Crosstalk between flavonoids and the clock is evident in photosynthetic rhythmicity

To determine whether the effects of flavonoids on clock gene expression translated to physiological outputs, we assessed the cycling of photosynthetic activity, an established indicator of endogenous clock function (Dodd et al., 2015; Shor et al., 2017; Guadagno et al., 2018). For this analysis, we first used hyperspectral imaging, which has been shown to provide an effective real-time, nondestructive measure of photosynthetic activity in individual soybean and wheat leaves (Pan et al., 2015). To test this in Arabidopsis, Col-0 and *tt4-11* plants were grown on soil for 5-6 weeks in 12h/12h LD, then transferred to LL and leaf reflectance measured from 385 to 1027 nm. This analysis uncovered a strong diurnal periodicity in three of five Col-0 plants examined over a 48h period at both 659 nm (Figure 4A) and 469 nm (Figure S2), the optimal wavelengths for detecting chlorophyll a and b. In contrast, no periodicity was observed in any of the five *tt4-11* plants that were analyzed.

**Figure 4.**
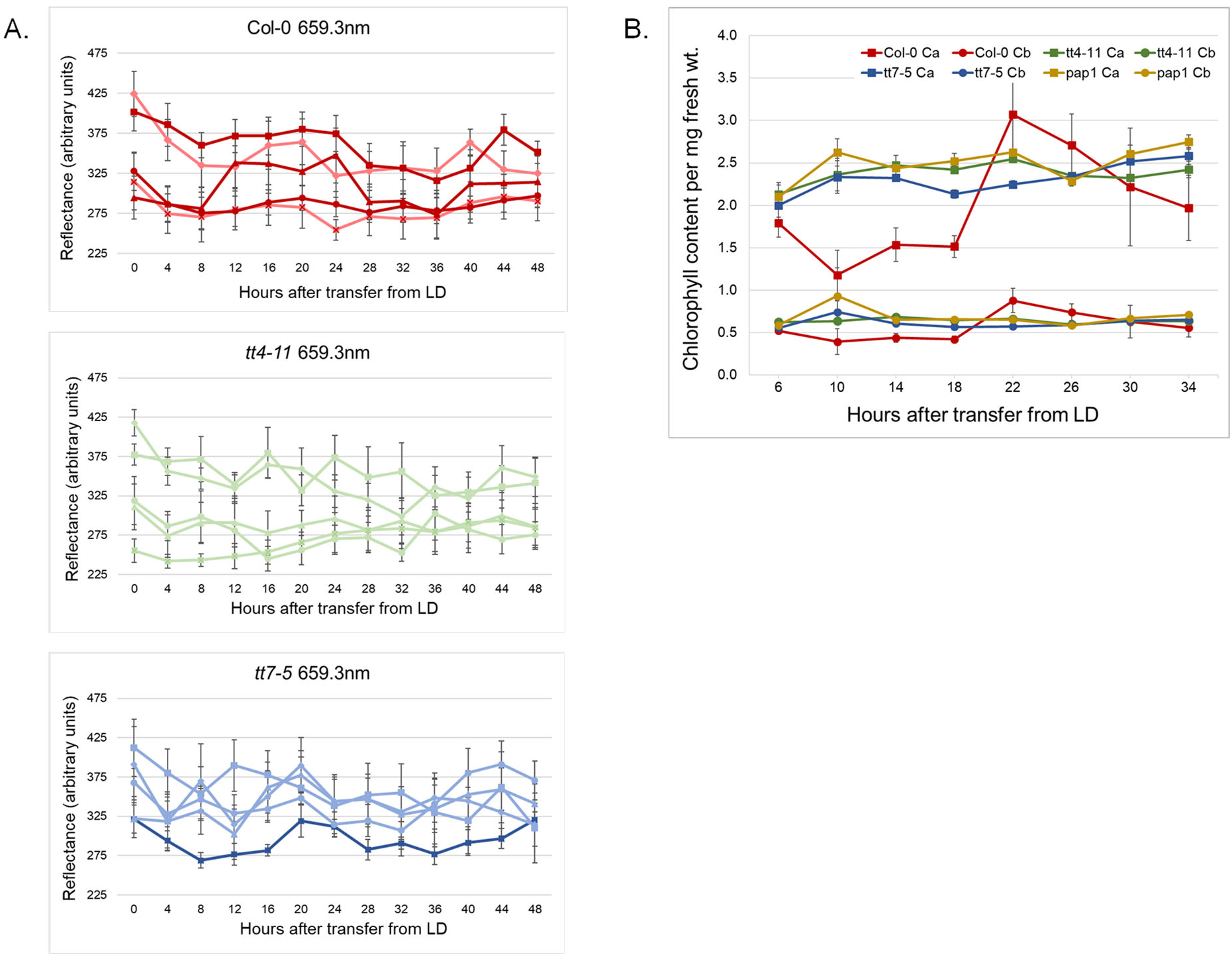
Effects of flavonoid mutations on cycling of chlorophyll levels in Arabidopsis plants. Plants were entrained in 12h/12h LD and transferred to LL before data or sample collection. (A) Hyperspectral imaging at the optimal wavelength for chlorophyll a (470 nm) and b (660 nm); lines represent the average reflectance of nine leaves from each of five 5-6 week-old plants tracked for 48h after shifting from LD to LL. Nine leaves were analyzed on each of five plants for each genotype. Dark lines represent plants exhibiting diurnal rhythmicity with p <0.05 per Metacycle analysis. (B) Spectrophotometric quantification of intracellular chlorophyll a and b content in above-ground parts of 4.5-week-old plants. Data points represent the average value calculated from 3-4 plants at each time point. Error bars in all panels are standard error.

This distinction was even more evident in a spectrophotometric analysis of chlorophyll content in 4.5-week-old plants. Using this more conventional approach (Barnes et al., 1992), *tt4-11* plants exhibited no evidence of cycling of chlorophyll a and b levels in contrast to the strong periodicity apparent in Col-0 plants (Figure 4B). These differences show that disruption of core clock and clock-regulated genes associated with the loss of CHS and/or flavonoids is reflected in alterations in clock-controlled physiology in these plants.

### Disruption of the dihydroxyflavonoid branch pathway alone also affects clock gene expression

To determine whether the loss of CHS protein or specific flavonoids are responsible for the observed effects on clock gene expression, we examined a null allele for *tt7,* the locus for the single-copy F3’H gene in Arabidopsis. Disruption of this gene affects one half of the central pathway, disabling the synthesis of quercetin, one of two abundant and ubiquitous flavonols present in Arabidopsis, as well as cyanidin and epicatechin, the primary anthocyanin and proanthocyanidin pigments, respectively, present in this species (Figure 1A). Analysis of the F3’H null allele, *tt7-5* (Bowerman et al., 2012), uncovered the same alterations in *CCA1* and *TOC1* promoter activity as we had observed for *tt4-11* (Figure 3). Effects on steady-state mRNA levels were also similar, even though these were again not all statistically significant (Figure 2). Importantly, *tt7-5* plants also lacked photosynthetic rhythmicity (Figure 4) Together, these results indicate that it is not lack of CHS protein, but the pathway end products, and dihydroxylated flavonoids such as quercetin and cyanidin, in particular, that are responsible for the observed effects of the *tt* mutants on clock function.

To examine this possibility further, we asked whether supplying flavonoids in the growth medium would alter *CCA1p::LUC* expression in wild-type seedlings. Previous work has established good evidence for the uptake and transport of exogenous flavonoids, including quercetin, from the growth medium by Arabidopsis plants (Buer et al., 2007). In this experiment, entrainment of Col-0 in 12h/12h LD on MS-sucrose medium containing 1 to 25 μM quercetin altered *CCA1* promoter activity in a dose-dependent manner, with low concentrations suppressing, and higher concentrations elevating promoter activity (Figure 5). These findings suggest both that quercetin is an active player in modulating rhythmicity in Arabidopsis and that there is an optimal intracellular level of flavonoids required for this effect. However, other dihydroxyflavonoids may have a similar effect, as growth on nobiletin, a flavonoid present at high levels in citrus but not produced by Arabidopsis, modified expression of the *CCA1* promoter in Col-0 seedlings, as did growth on the quercetin glycoside, rutin, and the flavonoid precursor, naringenin (Figure S2). These experiments also support the conclusion that it is flavonoid end products rather than the CHS enzyme that are responsible for the observed effects on circadian rhythmicity. This hypothesis is further supported by the finding that *pap1* seedlings, which accumulate high endogenous levels of flavonoids due to ectopic overexpression of a myb transcription factor (Borevitz et al., 2000), also lack rhythmic chlorophyll levels, similar to *tt4-1,* (Figure 4B). Together, these results further suggest that the antioxidant potential of flavonoids is not the only, or even the key, feature underlying the effects of flavonoids on clock activity.

**Figure 5.**
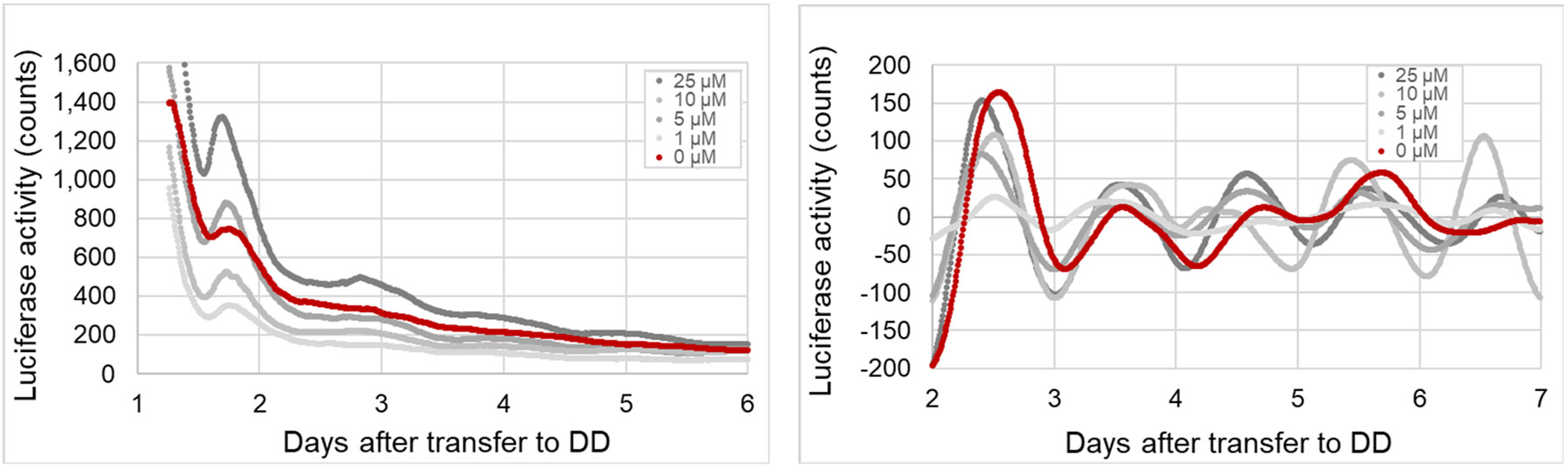
*CCA1p::LUC* expression in seedlings grown in the presence of different concentrations of quercetin. Luciferase activity was assessed as described for Figure 3 after 6d growth in 12h/12h LD. Data were normalized on a per-seedling basis, with 2-9 seedlings on each plate, with five plates per condition. Unprocessed (left) and detrended (right) data are shown.

## Discussion

Virtually every living organism possesses a clock mechanism that coordinates its biological processes with the rhythmic day/night cycle. Interestingly, the clock machinery has evolved independently in the different eukaryotic lineages so that the plant, insect, and animal clocks have few components in common, despite being organized around similar design principles. The plant clock machinery is particularly complex, reflecting its integration with response pathways for a wide range of environmental signals, from light and temperature to biotic and abiotic stress (Bendix et al., 2015; Creux and Harmer, 2019; McClung, 2019). Perhaps not surprisingly, the clock contributes to both plant resilience and productivity, and selection of circadian gene variants appears to have been key to the domestication of agriculturally-important plant species, affecting both flowering time and adaptation to stress (Creux and Harmer, 2019). As in all eukaryotes, the core clock is characterized by an interlocked transcription-translation feedback loop (TTFL), which in plants is centered on CCA1 and TOC1 as core components of the morning and evening clocks. The TTFL orchestrates the expression of a vast array of downstream genes and is itself regulated by a growing list of small molecules and hormones at both the transcriptional and post-transcriptional levels (Hearn and Webb, 2020). Signaling by reactive oxygen species (ROS) is emerging as a key element in a feedback system that coordinates the core circadian machinery with a wide variety of rhythmic cellular processes, including photosynthesis in plants (Guadagno et al., 2018; Simon et al., 2019).

Global analysis of gene expression in Arabidopsis *tt4* mutants has uncovered new evidence for crosstalk between the plant circadian clock and the abundant and ubiquitous phytochemicals known as flavonoids. Transcriptome, qPCR, and *in vivo* promoter analyses in flavonoid-deficient *tt4* seedlings indicate that clock gene expression is altered by disruption of the CHS locus. Similar observations for the F3’H null allele, *tt7-5,* suggest that dihydroxy B-ring flavonoids are a contributing factor. These conclusions are further supported by the finding that *tt4-11* and *tt7-5* plants exhibit marked defects in the periodicity of photosynthetic activity, the model readout of endogenous circadian function, and that growth on quercetin-containing medium affects expression of *CCA1p::LUC* differentially, depending on concentration, in wild-type seedlings. Interestingly, the WD-repeat protein, TTG1, was recently shown to be capable of mediating the plant circadian clock (Airoldi et al., 2019). This unique transcriptional regulator controls not only trichome development and the production of seed coat mucilage, but also the synthesis of anthocyanins and proanthocyanidins (Figure 1A). However, the effect of TTG1 on the plant clock has been attributed, not to any of these activities, but rather to retention of functionality present in a common ancestor of TTG1, and the related LIGHT-REGULATED WD1 and LIGHT-REGULATED WD2 clock regulators. This suggests that the molecules responsible for the observed crosstalk between flavonoids and the clock in *tt4-11* and *tt7-5* may well be products of early (more primitive) stages of the pathway leading to flavonols, at least in Arabidopsis.

Prior to the current study, there was just one publicly-available transcriptome dataset for a flavonoid-deficient plant line, used to examine the contribution of flavonoids to oxidative and drought tolerance (Nakabayashi et al., 2014). These experiments used a different allele of *tt4* (*tt4-3*, which is also a null allele in a Columbia background; Shikazono et al., 1998) and mature plants grown on 1% sucrose under 16h/8h LD conditions (versus 6-day-old seedlings, 2% sucrose, LL in our case). Our re-examination of this dataset using Genevestigator (Hruz et al., 2008) identified numerous core clock and clock-associated DEGs, echoing the results of our study, although consistently in the opposite direction (Dataset S5), similar to what we observed by qPCR in LD-entrained seedlings (Figure 2). Other genes unrelated to clock function did not exhibit this inverse relationship (Dataset S5). This provides further support for the finding that flavonoids contribute differentially to circadian rhythmicity under LL and LD conditions. It also underscores our finding that flavonoids influence the clock at different stages of development.

Despite decades of research on the physiological effects of flavonoids in plants, understanding of the underlying modes of action remains remarkably murky (Winkel, 2019; Agati et al., 2020). Flavonoid bioactivity has invariably been ascribed to the strong antioxidant potential of these compounds, which can be present at quite high levels in certain tissues and in response to diverse biotic and abiotic stresses. This may well be the case in some situations, for example, where flavonoids protect against oxidative stress (Nakabayashi et al., 2014; Kurepa et al., 2016; Muhlemann et al., 2018). There is also a strong connection between the clock and the intracellular redox state, which is modulated both by endogenous metabolic processes, including respiration and photosynthesis, and by environmental conditions that elevate intracellular ROS (Guadagno et al., 2018). It is therefore reasonable to suppose that flavonoids may contribute to maintaining ROS homeostasis and the integrity of the plant clock; a reduction in antioxidant potential could lie behind the observed disruption of clock gene expression and cycling of chlorophyll levels in the *tt4* and *tt7* lines.

One prediction of this hypothesis is that the transcriptomes of flavonoid mutant lines should display the hallmarks of oxidative stress under the growth conditions where altered circadian gene expression was observed. To examine this possibility we compared the high-confidence DEGs for *tt4-11* with the “common stress transcriptome,” 197 genes identified by analysis of publicly-available microarray data as being induced by a wide variety of biotic and abiotic stressors (Ma and Bohnert, 2007). This analysis showed that 25 of the 120 high-confidence upregulated genes in the *tt4-11* transcriptome were members of this group (Dataset S2), including the gene that was most highly induced, At1g74450. Although not yet functionally characterized, this gene has been shown to provide a link between multiple environmental stresses and plant resilience, including pollen development (Visscher et al., 2015). Overall, the *tt4-11* expression profile suggests that the absence of flavonoids does indeed result in a persistent, if modest, stress condition in LL-grown seedlings; it remains to be determined if this is also the case in LD.

To examine whether this response related specifically to a lack of antioxidant potential in *tt4-11*, the high-confidence list was further compared to the genes shown by Gadjev et al. (2006) to exhibit altered expression in ROS-sensitive Arabidopsis lines. However, at most 3 to 4 of the multiple genes defined in mutant lines deficient in the response to singlet oxygen, superoxide, or H_2_O_2_ were also mis-expressed in *tt4-11,* and in many cases showed the opposite change, being induced rather than suppressed and *vice versa* (Dataset S2). The *tt4-11* DEGs did include the map kinase, MAPKKK14 (log2fold 1.329 = 2.86-fold), but no other components of the stress-response associated mitogen-activated kinase (MAPK) pathway. There was also modest elevation (log2fold 0.713 = 1.64-fold) of AGC2, a protein that is strongly induced by singlet oxygen, the main ROS produced in plants under high light associated with photooxidative damage in chloroplasts under excess light energy (Shumbe et al., 2016). In addition, the *CAT2* gene, which encodes the predominant catalase critical for scavenging H_2_O_2_ in Arabidopsis (Mhamdi et al., 2010), exhibited elevated expression in the *tt4-11* LL transcriptome (log2fold 1.036 = 2.61-fold). However, this enzyme is also tightly linked with control of the central clock machinery in plants, as well as other organisms (Portolés and Más, 2010). Moreover, *AOX1a*, which encodes mitochondrial alternative oxidase, an important indicator of the response to excessive ROS, was unchanged and only two peroxiredoxin/like genes exhibited small changes (one up and one down) in expression, and neither has been linked to circadian sensing (Lee et al., 2018).

Overall, the *tt4-11* DEG dataset gave little support for the hypothesis that these fully flavonoid-deficient seedlings were suffering from a lack of antioxidant capacity. This is in spite of good evidence that the converse situation, overaccumulation of flavonoids in mutant lines or by feeding, can protect plants from oxidative stress (Nakabayashi et al., 2014; Kurepa et al., 2016). This is also evidenced by the finding that *pap1* seedlings, with severely-elevated endogenous levels of flavonoids, exhibited similarly arrhythmic chlorophyll levels as the flavonoid-deficient lines (Figure 4). Gadjev found that, although transcripts involved in flavonoid biosynthesis are induced in response to singlet oxygen, these are downregulated in response to H_2_O_2_, consistent with evidence that catalases and peroxidases are substantially inhibited by flavonoids (Gebicka, 2020).

In fact, there has been substantial debate regarding the role of flavonoids as antioxidants in plants, including from the viewpoint that intracellular concentrations vary tremendously across compartments, with the majority of flavonoids localized within the vacuole, the cell wall, and in at least some cases the nucleus (Winkel, 2019; Agati et al., 2020). There has been little exploration of alternative explanations for flavonoid action in plants, such as modulation of the activities of enzymes or signaling molecules, despite long-standing evidence of their potent and specific protein-binding potential (Gebicka, 2020). Similarly, in animals it has been argued that, other than in red blood cells and the intestine, concentrations are far too low for flavonoids to serve as antioxidants to provide protection against ROS. Instead, the focus of efforts to understand flavonoid function in animals is increasingly on interaction with components of intracellular signaling cascades as well as metabolic enzymes and drug transporters, both *in vitro* and *in vivo* (Middleton et al., 2000; Crozier et al., 2009; Miron et al., 2017; Gebicka, 2020).

Interestingly, it has recently been shown that a number of different flavonoids can modulate various aspects of the circadian clock in cultured animal cells and in mice (Xu and Lu, 2019). These studies provided evidence that alteration of circadian cycling by the polymethoxy flavone, nobiletin, involves binding to components of the ERK/CREB and REV-ERB/ROR signaling pathways (He et al., 2016; Shinozaki et al., 2017). Although homologous pathways are not present in plants, the component proteins are members of highly-conserved families that occur in all eukaryotes, suggesting that common mechanisms of action may exist, even if these involve different specific targets. Identification of protein-binding targets of flavonoids in plants and other organisms lags well behind animals. However, there are a few notable examples, including direct binding and inhibition by quercetin of the plant-specific protein serine/threonine kinase, PINOID, proposed to be the mechanism by which flavonoids control polar auxin transport (Henrichs et al., 2012). Defining the mechanistic basis of flavonoid action in plants through interaction with proteins and other biomolecules remains a fruitful area for further exploration.

The evidence for crosstalk between flavonoids and circadian cycling is a new and intriguing example of the diverse and potent biochemical activities of plant specialized metabolites, both in plants and the organisms that consume them. Although the current study represents the first report for this association in plant systems, Hu et al. (2020) recently noted enhanced expression of the clock-associated genes, *PRR5*, *FT,* and *LHY*, in two flavonoid hyper-accumulating lines of licorice, which suggests that the connection between flavonoids and the plant clock may well be a widespread phenomenon. The similar effects in animals, despite the dissimilarity of the clock machinery, raises the question of whether flavonoids also influence clock activity in yet other organisms, such as insects and microbes, and opens new doors to understanding the mode(s) of action of these potent and ubiquitous natural products in an important new context.

## Materials and Methods

### Plant genotypes and growth conditions

Flavonoid mutant lines, *tt4-2* (back-crossed to wild type to remove a *max1* mutation in the background), *tt4-11* (SALK_020583), and *tt7-5* (SALK_053394), were described previously (Burbulis et al., 1996; Bennett et al., 2006; Lewis et al., 2011; Bowerman et al., 2012). Comparisons were made against Col-0 (CS7000) for *tt4-11* and *tt7-5,* or against a Columbia wild-type line originally obtained in Howard Goodman’s laboratory at Harvard Medical School for *tt4-2*. The *CCA1p::LUC* and *TOC1p::LUC* promoter reporter lines were a generous gift of Rob McClung (Dartmouth College). These were crossed with *tt4-11* and *tt7-5* using the mutant lines as the maternal parents, and F2 plants homozygous for the flavonoid gene mutation were selected based on the *transparent testa* phenotype. Multiple independent F3 lines were used for measurements of circadian parameters.

### RNA-Seq and qRT-PCR

Gene expression analyses were performed using five-day-old whole seedlings. Seeds were sown on 1X MS medium, pH 5.7, 2% sucrose, 0.8% agar as described previously (Kubasek et al., 1992), in 8 or 15 mm Petri dishes. Following stratification at 4°C in darkness for 3-5 days, seedlings were grown at 22°C in LL or a 12h/12h LD cycle (~100 μE). At 5 days following germination, seedlings were harvested and weighed, then in 100 mg aliquots, flash frozen in liquid nitrogen, and stored at −80°C prior to processing. For RNA-Seq analysis, samples were collected as 100 mg aliquots within a one hour time span, although the precise time of day was not recorded.

RNA was extracted using the Qiagen RNeasy Plant Maxi or Mini Kit with DNase on-column digestion (Qiagen, Germantown, MD). For RNA-Seq analysis, libraries (four wild-type and four *tt4-11*) were prepared at the Virginia Tech Genomics Core Sequencing Laboratory using the TruSeq Stranded mRNA HT Sample Prep Kit (Illumina), which uses polyA enrichment. These were sequenced on a HiSeq 2500 (Illumina) using Rapid Run 1×100 single read cycle clustering and sequencing. The resulting data were processed by the Virginia Tech Advanced Research Computing (ARC) core facility as described in Chen et al. (2016). One of the *tt4-11* libraries was eliminated from the final analysis. Genes found to meet the cutoff for differential expression in *tt4-11* and wild-type (log2fold of |±1.0| = ~2.2 fold, all having p values <<0.001) were categorized based on function using the Bio-analytical Resource for Plant Biology’s Classification SuperViewer (http://bar.utoronto.ca/). Genes related to clock function were curated manually. Microarray data (dataset AT-00697) described in Nakabayashi et al. (2014) was analyzed using Genevestigator (Hruz et al., 2008).

For qRT-PCR analysis, cDNA synthesis was carried out using SuperScript IV reverse transcriptase and RNaseOUT ribonuclease inhibitor (both from Invitrogen) and 2 μg of total RNA for three biological replicates of each genotype (*tt4-2, tt4-11,* and *tt7-5*, and wild type). qRT-PCR was performed using *Power* SYBR Green PCR Master Mix (Life Technologies Corp.) using the primer pairs listed in Table S3. Analyses were carried out in 384-well plates in a QuantStudio 6Flex (Applied Biosystems) with three technical replicates per sample. The resulting data were analyzed using the ∆∆C_T_ method. Statistical analysis was performed using JMP Pro 15.0.0. (SAS Institute, Inc.).

### Real-time bioluminescence measurements of promoter activity

Luciferase bioluminescence measurements were performed using seedlings containing *CCA1p::LUC* or *TOC1p::LUC* constructs. Growth was on MS-sucrose-agar medium as described above, but using 35 mm dishes containing equivalent numbers (typically 5-15) of seedlings per dish. Medium for quercetin feeding experiments was prepared using a 100 mM stock solution in ethanol. Following stratification for 2-3 days at 4°C in darkness, seedlings were germinated under a 12 h/12h LD cycle or continuous light for 5 days. A 100 μl aliquot of 1 mM luciferin was added to the surface of each plate and reporter activity was recorded using a LumiCycle luminometer (Actimetrics, Inc.) for up to 10 days. Detrended data were fit to a polynomial of 3 and smoothed over 30 min intervals using ClockLab (Actimetrics, Inc.) and amplitude, phase, and period quantified using FFT NLLS analysis in BioDare2 (Zielinski et al., 2014). Each experiment was repeated independently at least once.

### Analysis of photosynthetic activity

Analyses were carried out on mature (5-6 week-old) soil-grown plants. Col-0, *tt4-11*, and *tt7-5* plants were entrained to a 12h/12h LD cycle, and moved to continuous light on the first day of data collection. Plants were imaged starting at Circadian Time (CT) 6 every 4 hours for 48 hours. The hyperspectral imaging system consisted of a Pika L hyperspectral imaging camera (385.6– 1027 nm spectral range with 2.0 nm resolution), linear translation stage, mounting tower, lighting assembly, and a desktop computer loaded with Spectronon-Pro software to control the imager and translation stage during imaging (Resonon, Bozeman, MT). This software was also used to measure reflectance values from nine leaves per plant and the average used to calculate the reflectance values per plant per time point. Statistical analysis of rhythmicity was performed using meta_2d p-value in MetaCycle, combining algorithms JTK_CYCLE, ARSER (ARS), and Lomb-Scargle (LS) (Wu et al., 2015). Spectrophotometric quantification of chlorophyll content was carried largely as described in Yoo et al. (2019). Briefly, 4.5-week-old plants were transferred to LL and above-ground tissues from individual plants were harvested, weighed, and quick-frozen in liquid nitrogen, then stored at −80°C prior to processing. Chlorophyll was extracted in 1.5 ml DMSO at 65°C for 1h. Absorbance of 50 μl aliquots was determined at 648 and 665 nm using a Cytation 5 plate reader (BioTek, Winooski, VT) and content of chlorophyll a and b per μg wet weight determined using the equations in Barnes et al. (1992).

## Supporting materials

**Figure S1:**
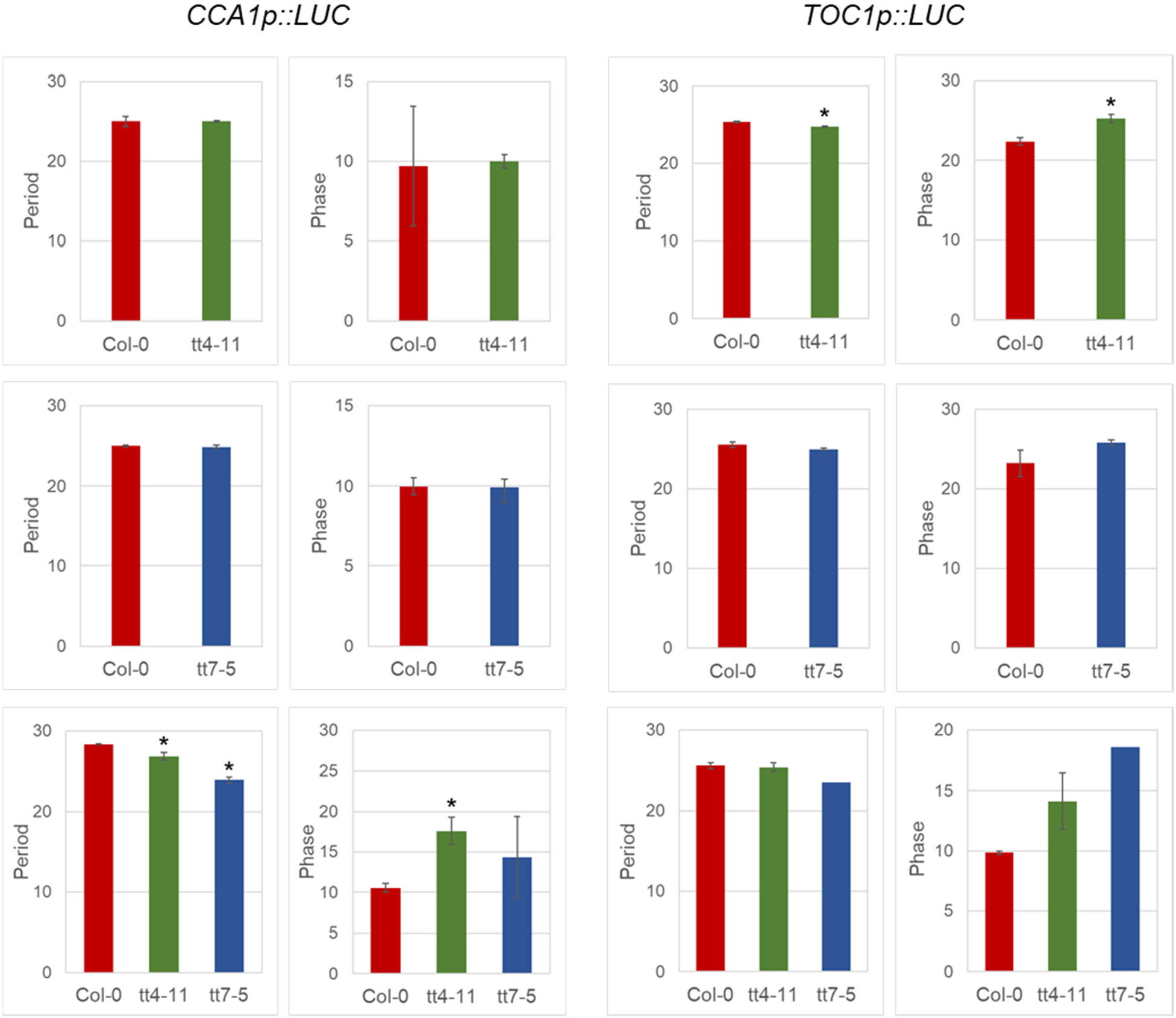
Period and phase of *CCA1p::LUC* and *TOC1p::LUC* expression in *tt4-11* and *tt7-5* seedlings. Luciferase activity was assessed in seedlings using a Lumicycle following transfer from 12h LD (top two rows) or LL (bottom row) to continuous darkness (DD). In all panels, results are shown for Col-0 (red), *tt4-11* (green), and *tt7-5* (blue) seedlings. Each line represents the average of four biological replicates for Col-0 and the average of 2-4 replicates for each of three independent F3 lines for *tt4-11* and *tt7-5*. Error bars are standard error; * indicates Student’s t-test p-value <0.05.

**Figure S2:**
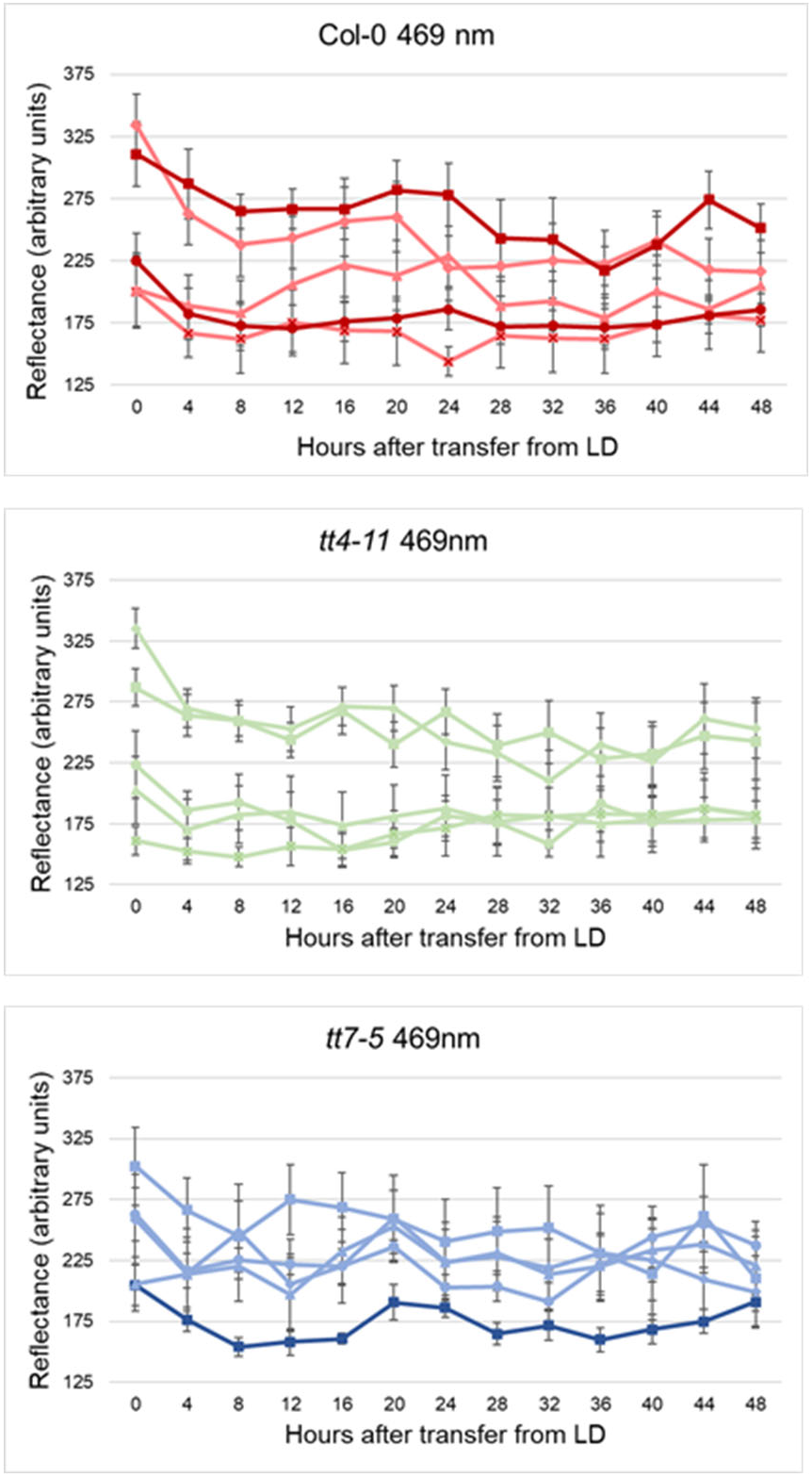
Hyperspectral imaging analysis of leaves from mature plants over 48h time course at 469 nm. Plants were entrained in 12h/12h LD and transferred to LL before data or sample collection. Lines represent the average reflectance of nine leaves from each of five 5-6 week-old plants tracked for 48h after shifting from LD to LL. Nine leaves were analyzed on each of five plants for each genotype. Dark lines represent plants exhibiting diurnal rhythmicity with p <0.05 per MetaCycle analysis. Error bars are standard error.

**Figure S3:**
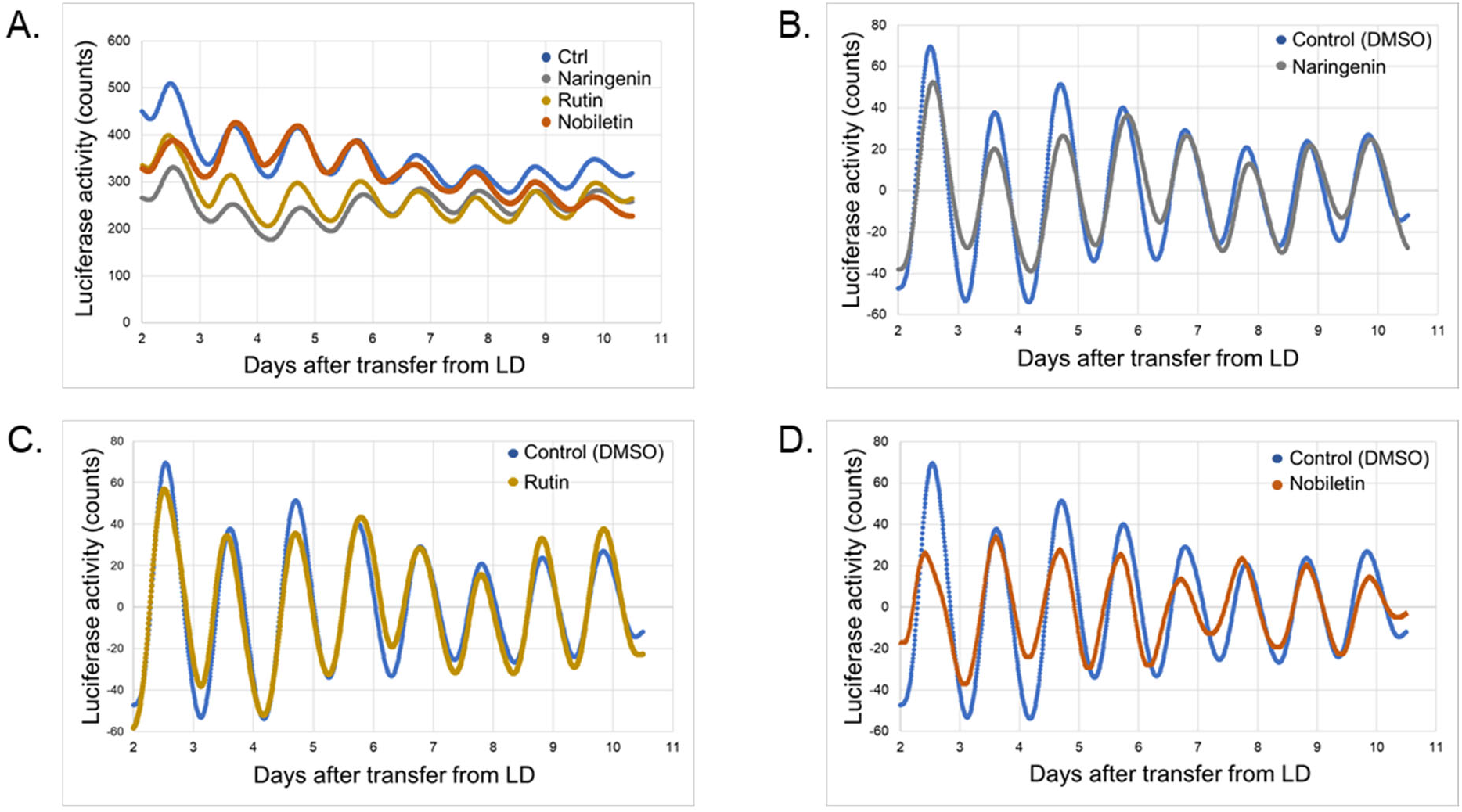
Effects of flavonoids on *CCA1p::LUC* rhythms in Col-0 (wild-type) seedlings. Luciferase activity was assessed in seedlings using a Lumicycle following transfer from 12h/12h LD to continuous darkness (DD). (A) Unprocessed traces and (B-D) detrended (baseline-subtracted) data comparing seedlings growing on medium containing no added flavonoids or with 100 μM naringenin, rutin, or nobiletin. Data were normalized on a per-seedling basis.

**Table S1:**
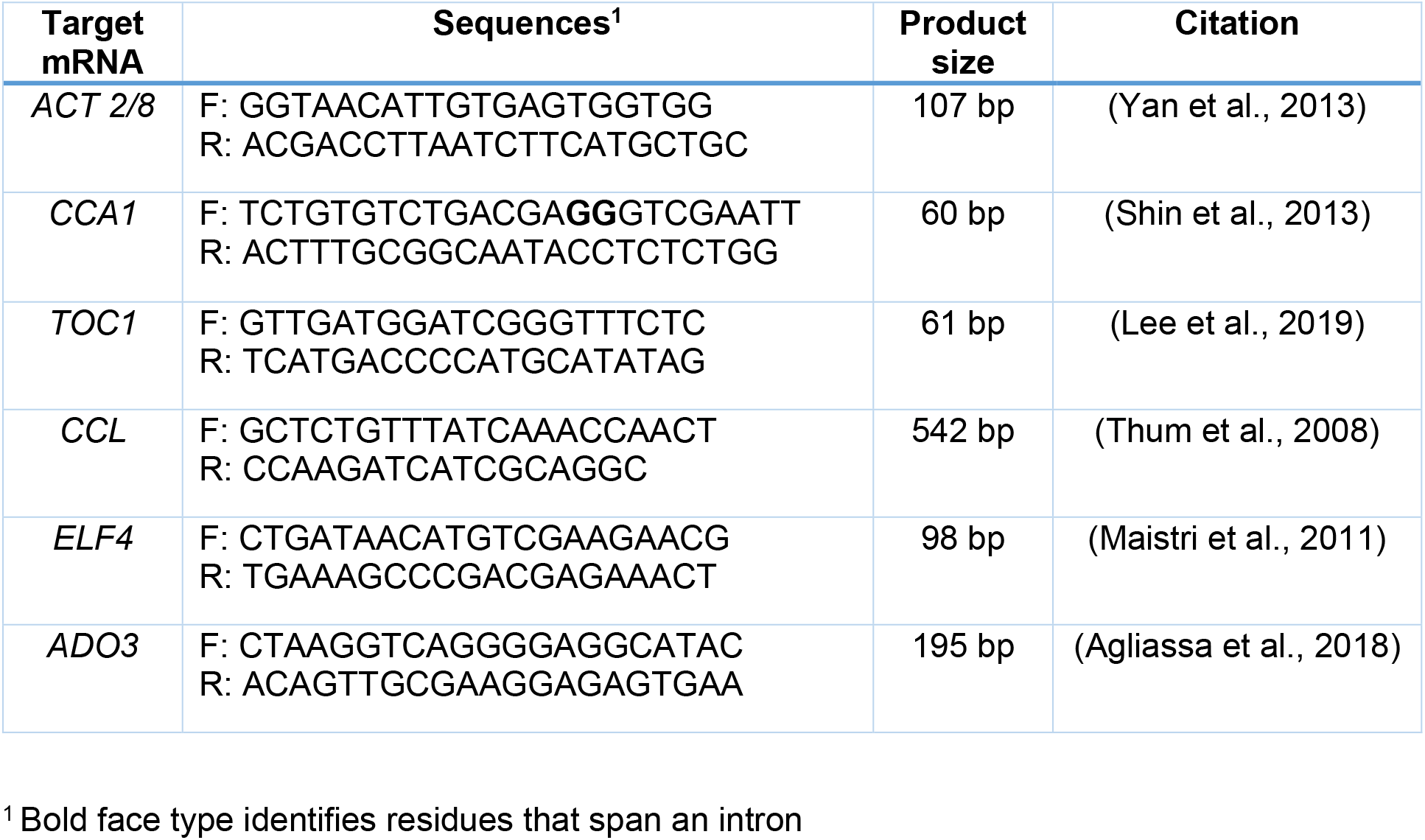
Primers used for qPCR

**Dataset S1**: Full RNA-Seq data with annotations

**Dataset S2**: High-confidence differentially-expressed genes (DEGs) (log2fold |±1.0|) and comparisons with previously-published lists of: 1) circadian-regulated genes in Arabidopsis (Covington et al., 2008); 2) genes with proximal *CCA1* binding sites (Nagel et al., 2015); 3) genes encompassing universal stress transcriptome (Ma and Bohnert, 2007); and 4) genes represented in reactive oxygen species (ROS)-responsive transcriptomes (Gadjev et al., 2006).

**Dataset S3**: DEGs with log2fold≤|0.50| & p<.05

**Dataset S4**: Panther overrepresentation analysis

**Dataset S5**: Comparison of RNA-Seq and previously-published microarray results

## Acknowledgments

This work was supported by a grant from the National Science Foundation (grant IOB-0820674 to B.S.J.W.), funding from the Federal Work-Study Program and a Fralin Life Sciences Institute/Translational Plant Sciences Summer Undergraduate Fellowship (to L.C.C.), and financial support from the College of Science and the Department of Biological Sciences at Virginia Tech. The authors thank John McDowell for providing Col-0 (CS7000) seeds, Gloria Muday (Wake Forest University) for the *tt4-2* back-crossed line, and Rob McClung (Dartmouth College) for the wild-type *CCA1p::LUC* and *TOC1p::LUC* lines. We thank Saikumar Karyala of the Genomics Sequencing Center for expert advice on the RNA-Seq experiment, Robert Settlage of Advanced Research Computing at Virginia Tech and for the initial bioinformatics analysis, and Song Li and Kshitiz Dhakal for providing resources and expertise for hyperspectral imaging analysis.

## Author contributions

BSJW conceived the project. SBH, LCC, GCP, EL, RS, SK, and BSJW designed and performed experiments, analyzed the data, and contributed to the original draft of the manuscript. SBH, EL, SK, and BSJW revised and edited the manuscript; all authors read and approved the final version.

## Data availability

The data that support the findings of this study will be deposited in the NCBI Gene Expression Omnibus (GEO) archive at https://www.ncbi.nlm.nih.gov/geo/.

